# Tools for Genetic Engineering and Gene Expression Control in *Novosphingobium aromaticivorans* and *Rhodobacter sphaeroides*

**DOI:** 10.1101/2023.08.25.554875

**Authors:** Ashley N. Hall, Benjamin W. Hall, Kyle J. Kinney, Gabby G. Olsen, Amy B. Banta, Daniel R. Noguera, Timothy J. Donohue, Jason M. Peters

## Abstract

Alphaproteobacteria have a variety of cellular and metabolic features that provide important insights into biological systems and enable biotechnologies. For example, some species are capable of converting plant biomass into valuable biofuels and bioproducts have the potential to form the backbone of the sustainable bioeconomy. Among the Alphaproteobacteria, *Novosphingobium aromaticivorans*, *Rhodobacter sphaeroides*, and *Zymomonas mobilis*, show particular promise as organisms that can be engineered to convert extracted plant lignin or sugars into bioproducts and biofuels. Genetic manipulation of these bacteria is needed to introduce engineered pathways and modulate expression of native genes with the goal of enhancing bioproduct output. Although recent work has expanded the genetic toolkit for *Z. mobilis*, *N. aromaticivorans* and *R. sphaeroides* still need facile, reliable approaches to deliver genetic payloads to the genome and to control gene expression. Here, we expand the platform of genetic tools for *N. aromaticivorans* and *R. sphaeroides* to address these issues. We demonstrate that Tn*7* transposition is an effective approach for introducing engineered DNA into the chromosome of *N. aromaticivorans* and *R. sphaeroides*. We screen a synthetic promoter library to identify inducible promoters with strong, regulated activity in both organisms. Combining Tn*7* integration with promoters from our library, we establish CRISPR interference systems for *N. aromaticivorans* and *R. sphaeroides* that can target essential genes and modulate engineered pathways. We anticipate that these systems will greatly facilitate both genetic engineering and gene function discovery efforts in these industrially important species and other Alphaproteobacteria.

**IMPORTANCE:** It is important to increase our understanding of the microbial world to improve health, agriculture, the environment and biotechnology. For example, building a sustainable bioeconomy depends on the efficient conversion of plant material to valuable biofuels and bioproducts by microbes. One limitation in this conversion process is that microbes with otherwise excellent properties for conversion are challenging to genetically engineer. Here, we report systems to overcome that barrier in the Alphaproteobacteria, *Novosphingobium aromaticivorans* and *Rhodobacter sphaeroides*, by producing genetic tools that allow easy insertion of engineered pathways into their genomes and to precisely control gene expression by inducing genes with synthetic promoters or repressing genes using CRISPR interference. These tools can be used in future work to gain additional insight into these and other Alphaproteobacteria and to optimize yield of biofuels and bioproducts.

## INTRODUCTION

A myriad of microbial activities has and can have positive impacts on the health of the planet, its inhabitants, and the production of compounds needed by a growing population. In recent years a diverse group of microbes have been identified that can convert renewable plant material into sustainable biofuels and bioproducts. However, additional genetic tools for these organisms are needed to dissect their metabolic and regulatory networks and to build economically feasible production strains. To generate the knowledge needed to power the future bioeconomy, we need the ability to easily engineer the genomes of additional microbial chassis and to rationally control the expression of bioproduct-relevant genes.

Many Alphaproteobacteria have unique metabolic pathways that make them ideal chassis for production of biofuels and bioproducts. *Novosphingobium aromaticivorans* and *Rhodobacter sphaeroides* are two Alphaproteobacterial species of particular interest for converting plant feedstocks into bioproducts. *N. aromaticivorans* utilizes aromatic carbon sources and has been engineered to produce bioproducts including pimelic acid and 2-pyrone-4,6-dicarboxylic acid (PDC), a nylon precursor from plant phenolics (1–3). *R. sphaeroides* is a well-studied photosynthetic bacterium with the capacity to produce biotechnologically useful products such as hydrogen, terpenes, and polyhydroxyalkanoates (4–11). The genetic tools to manipulate these bacteria include homology-based genome integration, Tn-seq, and a few inducible plasmid vectors (12–14). While effective in specific use cases, additional genetic tools are needed to overcome limitations in homology-based integration, make it easier to target essential or other specific genes, or bypass the use of plasmids.

The ability to generate and control expression of engineered pathways can also be critical for optimizing or altering bioproduct yields. Genomic integration of engineered pathways allows for stable maintenance of the pathway without continuous application of selective pressure (e.g., antibiotics) that is typically required to maintain plasmids (15). Various approaches exist for genomic integration of engineered DNA including homologous recombination, phage integrase systems, and transposons (16–18). Homologous recombination has the advantage of targeting DNA to a specific locus. However, single crossover events are often unstable without selection and stable double-crossovers typically require the use of a counter-selectable marker to enrich for the second crossover event, making homologous recombination time consuming and low efficiency. Integrases require specific sites on the genome to insert their DNA payload which must first be engineered into the target strains. Recent work has shown that arrays of integrase sites can be delivered to recipient genomes and the utilized for multiple insertions of cargo (19). While these systems are valuable for iterative strain engineering, they require a two-step procedure to add the initial integrase sites. Transposon-based integration of DNA has been used in diverse bacteria and is a highly effective, one-step procedure that can be implemented at the library scale. Although many transposons integrate randomly, the Tn*7* transposon from *Escherichia coli* integrates downstream of a conserved recognition site in the 3’ end of the *glmS* gene (20). Because *glmS* is conserved in most bacteria and because Tn*7* transposase recognizes only non-wobble bases in the *glmS sequence*, the Tn*7* recognition site is present in a broad set of bacterial genomes (21, 22). As such, Tn*7* has become an integration tool of choice in diverse bacteria including the Alphaproteobacterium *Zymomonas mobilis* (20, 23, 24). However, Tn*7* function has neither been demonstrated for *N. aromaticivorans* nor optimized for *R. sphaeroides* (25).

Methods to control gene expression, including inducible promoters and CRISPR interference (CRISPRi) knockdowns are useful for modulating the levels and temporal dynamics of gene products. Most inducible promoters are based on the consensus sequence from *E. coli* σ^70^; however, it is unknown how well these elements function in high GC-Alphaproteobacteria, such as *N. aromaticivorans* (63% GC) and *R. sphaeroides* (68% GC). For instance, the wild-type *lac* promoter from *E. coli* is reported to function poorly in *R. sphaeroides* (26). While no inducible promoters have been examined in *N. aromaticivorans* to our knowledge, promoters inducible by light, oxygen, crystal violet, and IPTG have been tested in *R. sphaeroides* (26–29). Although these foreign promoters can respond to an array of possible inducers, some inducers inhibit *R. sphaeroides* growth (crystal violet) (26) or substantially alter physiological processes of interest (light, oxygen). Existing IPTG inducible promoters are derivatives of the 16S rRNA promoter (11, 26) which is regulated according to nutrient availability by several cellular factors including CarD, ppGpp, and DksA in Alphaproteobacteria and some other species (30–33). Synthetic, IPTG-inducible promoters have a potential advantage of not being inherently regulated by nutrient conditions and of utilizing an inducer that is likely less physiologically disruptive to the host.

The development of facile genetic engineering tools, such as CRISPRi, depends on reliable strategies for integration of CRISPR systems into genomic DNA and inducible expression of CRISPRi components. We previously developed a platform called “Mobile-CRISPRi” that takes advantage of Tn*7*- based integration and synthetic promoters to deliver and induce CRISPRi knockdown in a number of diverse bacteria (23). Developing an effective Mobile-CRISPRi system for *Z. mobilis* required optimization of CRISPRi component expression (sgRNAs and dCas9), a phenomenon we observed in some, but not all recipient bacteria (23, 24).

In this work, we demonstrate the use of Tn*7* for stable, site-specific integration in *R. sphaeroides* and *N. aromaticivorans*, optimizing transconjugant recovery using several mating schemes. We create a series of strong, IPTG-inducible promoters that function in both *N. aromaticivorans and R. sphaeroides*, providing a reliable system for controlled gene expression. Finally, we combine these elements to demonstrate effective CRISPRi control systems that can phenotype essential genes in both organisms and control expression of engineered pathways for synthesis of valuable, foreign carotenoids in *N. aromaticivorans*.

## RESULTS

### Tn*7* integrates engineered DNA into *N. aromaticivorans* and *R. sphaeroides*

Site-specific Tn*7* transposition is a common method to integrate engineered DNA into the genomes of diverse bacteria (20, 22). We previously used a tri-parental mating scheme to introduce Tn*7* into the genomes of various Gammaproteobacteria (23). In this scheme, a recipient strain was mated with two *E. coli* donors: one harboring a plasmid containing Tn*7* transposase genes and one harboring a plasmid containing the Tn*7* transposon (23, 24, 34). Both plasmids require an *E. coli* host containing the *pir*^+^ gene for replication (R6Kγ origin) and cannot replicate in the recipient. To test Tn*7* integration as a vehicle to engineer *N. aromaticivorans* and *R. sphaeroides*, we attempted this and related mating schemes (Fig. 1AB).

We successfully used bi-parental mating to introduce Tn*7* into the *N. aromaticivorans* genome, albeit with variable efficiency (Table S1, Fig S1). In these transposition experiments, the percentage of transconjugants obtained per viable cells ranged from approximately 0.001-0.01%, but the total number of transconjugants never exceeded 10,000 per mL. However, subsequent mating experiments that included a conjugal helper strain (*E. coli* containing pEVS104 (35, 36)) in a quad-parental mating increased Tn*7* transposition efficiency when used with the same *N. aromaticivorans* recipient culture under otherwise identical conditions. (Fig.1C). We confirmed that insertion of the Tn*7* transposon did not affect the growth of *N. aromaticivorans* in rich and minimal media (Fig. 1E). To gauge the stability of Tn*7* integrants, we sequentially cultured *N. aromaticivorans* cells containing a Tn*7* transposon for approximately 50 generations in the absence of selection. We found that 100% of cells maintained the Tn*7* selectable marker (Fig. 1G).

**Figure 1:**
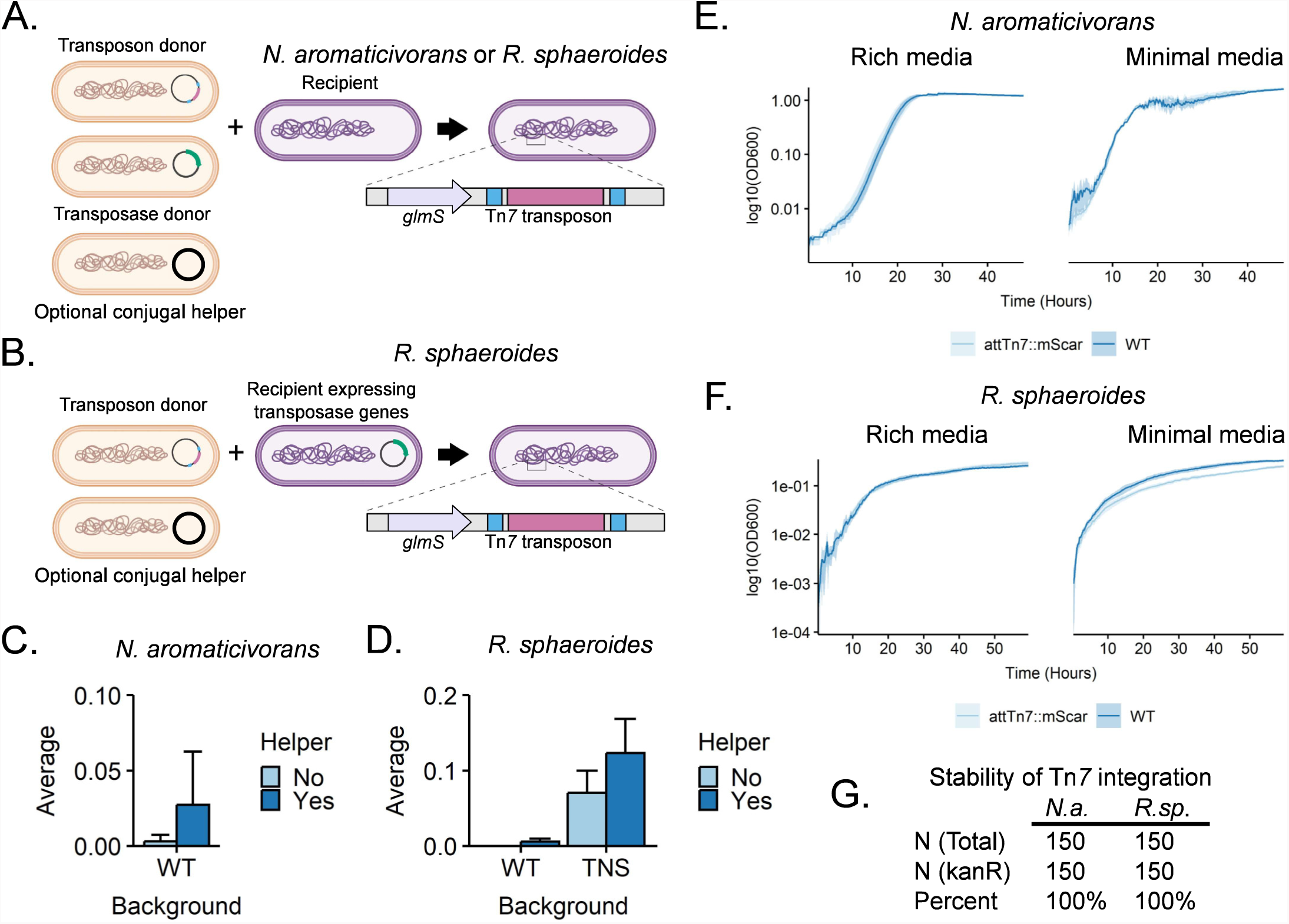
Insertion of Tn*7* into the genomes of *N. aromaticivorans* and *R. sphaeroides*. A: Schematic of a mating scheme for transfer of the Tn*7* transposon and Tn*7* transposase-expressing plasmid into the recipient of interest. B: Schematic of a mating scheme for delivery of a Tn*7* transposon into *R. sphaeroides* constitutively expressed with plasmid-encoded Tn*7* transposase genes (*tnsABCD*). C&D: Transposition efficiency into *N. aromaticivorans* and *R. sphaeroides,* respectively, using different mating schemes. E: Growth of *N. aromaticivorans* strains with or without att::Tn*7* insertions in rich (464a) and minimal (SIS+glucose) media. F: Growth of *R. sphaeroides* strains with or without att::Tn*7* insertions in rich (LB) and minimal (SIS) media. G: Stability of a *kanR* Tn*7* insertion in both *N. aromaticivorans* and *R. sphaeroides* following serial passaging in the absence of selection. *N.a.=N. aromaticivorans*, *R.sp.*=*R. sphaeroides*. Summary statistics and growth curve data can be found in tables S2-S5.

Tn*7* transposition into the *R. sphaeroides* genome at similar efficiencies required additional optimization. Our initial tri-parental mating scheme failed to recover a detectable number of transconjugants, and addition of the conjugal helper in a quad-parental mating resulted in fewer Tn*7* integrants than with *N. aromaticivorans* (Fig. 1D). We hypothesized that poor recovery of transconjugants was due to insufficient, transient expression of the Tn*7* transposase genes (*tnsABCD*). To test this hypothesis and increase Tn*7* transposition efficiencies, we cloned the transposase genes on a replicative plasmid in *R. sphaeroides* (pTNS++), producing a new recipient strain (Fig.1B). When using pTNS++ as a recipient, transposition efficiencies improved by more than an order of magnitude (from 0.006% to 0.12%) compared to cells lacking this plasmid (Fig. 1D). The pTNS++ plasmid is easily cured: 32% +/- 5% of colonies had lost the plasmid after a short period of growth in the absence of selection. In addition, we found that inclusion of the conjugal helper plasmid in the donor strain to a recipient that contains pTNS++ produced a modest increase in transconjugant recovery (less than 2-fold; Fig. 1D). We found that *R. sphaeroides* strains containing a Tn*7* insertion showed no growth defects in rich medium, but a small, statistically significant reduction of growth in minimal medium (generation time increases ∼1.6x, p< 0.05, Welch two-sample t-test) (Fig. 1F). As was the case with *N. aromaticivorans*, we also found that the Tn*7* insertion element remained stable when passaged in the absence of selection (Fig. 1G). From these experiments, we conclude that Tn*7* transposition is an effective and stable approach to integrate engineered DNA into *N. aromaticivorans* and *R. sphaeroides*. Below we describe some uses of this capability to gain important new information on these two alphaproteobacterial species.

### Synthetic, inducible promoters for *N. aromaticivorans* and *R. sphaeroides*

Synthetic, inducible promoters are valuable for strain engineering because they mitigate the effects of physiological regulation associated with native promoters and can be controlled in a temporal, reversible, and titratable manner. IPTG-inducible promoters are especially valuable for many studies because IPTG is metabolically inert, diffuses readily into most cells, and its interactions with Lac Repressor (LacI) are extremely well characterized (37–40). The native *lac* promoter from *E. coli* has been tested in *R. sphaeroides*, but expression was shown to be inconsistent and unreliable (26). Given the reported differences in promoter elements between alpahproteobacteria and other bacterial species (41) it is likely that the *E. coli lac* promoter will have limited utility in *N. aromaticivorans* and other organisms. To identify synthetic, regulated promoters for *N. aromaticivorans* and *R. sphaeroides*, we assembled a set of IPTG-inducible promoters (42) that vary in DNA sequences within the −35 and −10 elements (43), spacer length (44), presence or absence of an UP element (45), and *lac* operator sequence (46) (See Table 4). These promoters combined elements from the Anderson promoter library (47), UP sequences from Estrem et al. (48), and were based on a general understanding of *E. coli* σ^70^-DNA contacts (49–51).

To screen this promoter library for activity, we built a Tn*7*-based fluorescent reporter system. Each of the 20 promoters was independently cloned upstream of the *mScarlet-I* gene so their activities could be measured by mScarlet-I fluorescence. We integrated these constructs separately into the *N. aromaticivorans* and *R. sphaeroides att*_Tn*7*_ sites and into the *E. coli att*_Tn*7*_ site as a comparator (Fig.2B-D). Relative promoter activity was calculated by dividing the activity of each strain by the median activity of the least active of the 20 promoters in the respective host. For our analysis, synthetic promoters with the same or lower activity than a control strain lacking a promoter were considered to have negligible activity.

**Figure 2:**
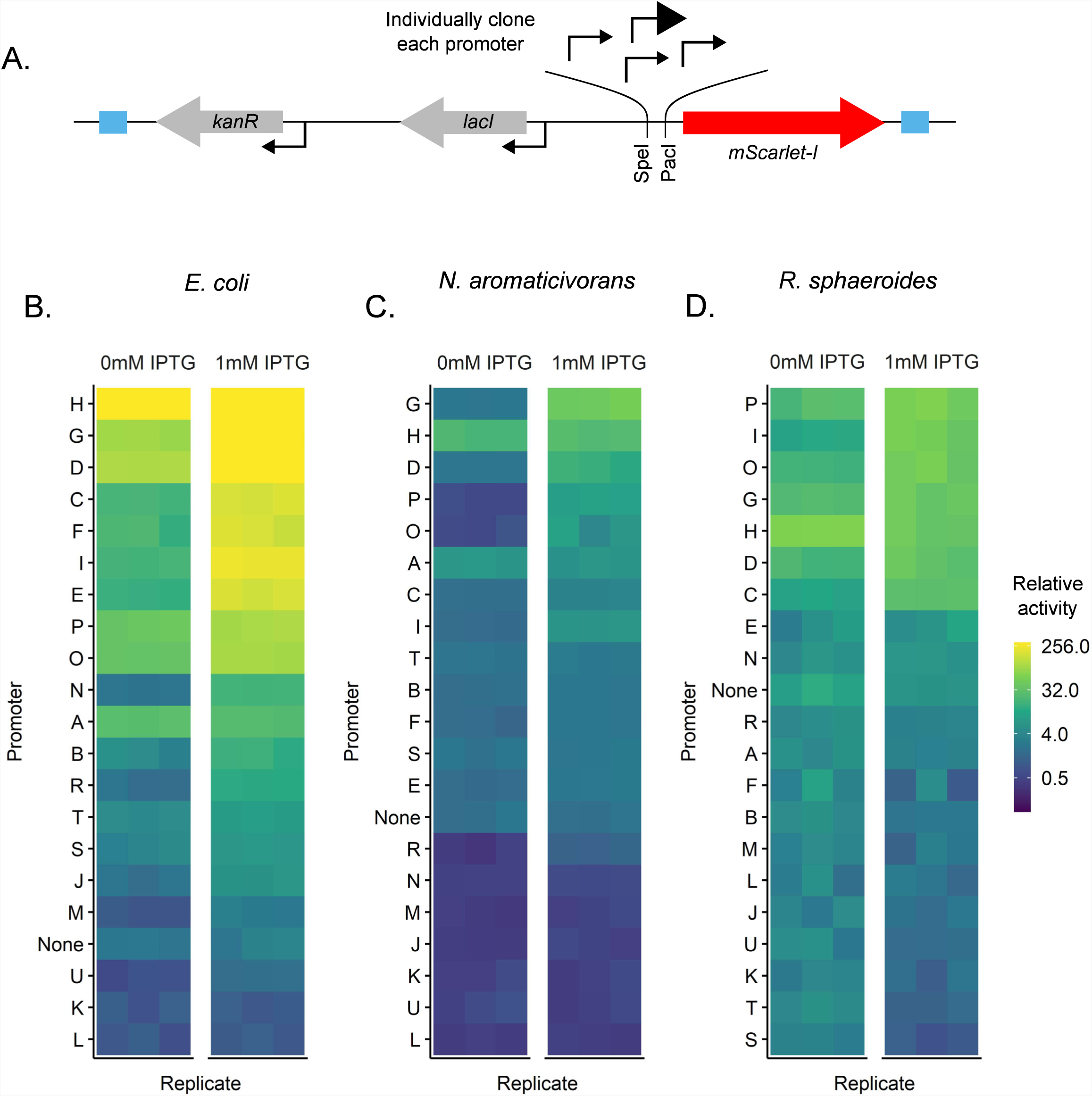
IPTG-inducible promoters expressing an *mScarlet-I* fluorescent reporter gene. A: Schematic of the Tn*7* transposon containing the test promoter construct (sequences of individual promoters provided in Table 4). Promoters of interest were cloned upstream of the *mScarlet-I* gene using the indicated PacI and SpeI restriction sites. B-D: Relative promoter activity in each *E. coli*, *N. aromaticivorans*, and *R. sphaeroides* in the presence and absence of IPTG. Relative activity was calculated by dividing each fluorescence-per-cell measurement by the median value of the least active promoter. Data are shown on a log2 scale. Promoter activity values and summary statistics can be found in tables S6-S14.

We found several synthetic promoters that had activity above background and were IPTG-inducible across the three organisms (Fig. 3A-B). In general, promoters D, G, O, and P (P_D_, P_G_, P_O_, P_P_) were active in both *N. aromaticivorans* and *R. sphaeroides*; these promoters contained near-consensus binding sites (−10 and −35) for *E. coli* σ^70^, underscoring the cross-species relevance of these sequences to activity. We also found that the most active *N. aromaticivorans* and *R. sphaeroides* promoters contained sequences with UP elements that had been previously isolated from a SELEX screen for tight binding to *E. coli* RNA polymerase (48). We found this later observation to be interesting because such AT-rich, “canonical” UP elements are essentially absent from known alphaproteobacterial promoters due to their high genomic GC content (65% and 68.5% GC for *N. aromaticivorans* and *R. sphaeroides*, respectively) (41). In summary, we have developed synthetic, IPTG-inducible promoters for *N. aromaticivorans* and *R. sphaeroides* that can be used to control expression of engineered genes or pathways.

**Figure 3:**
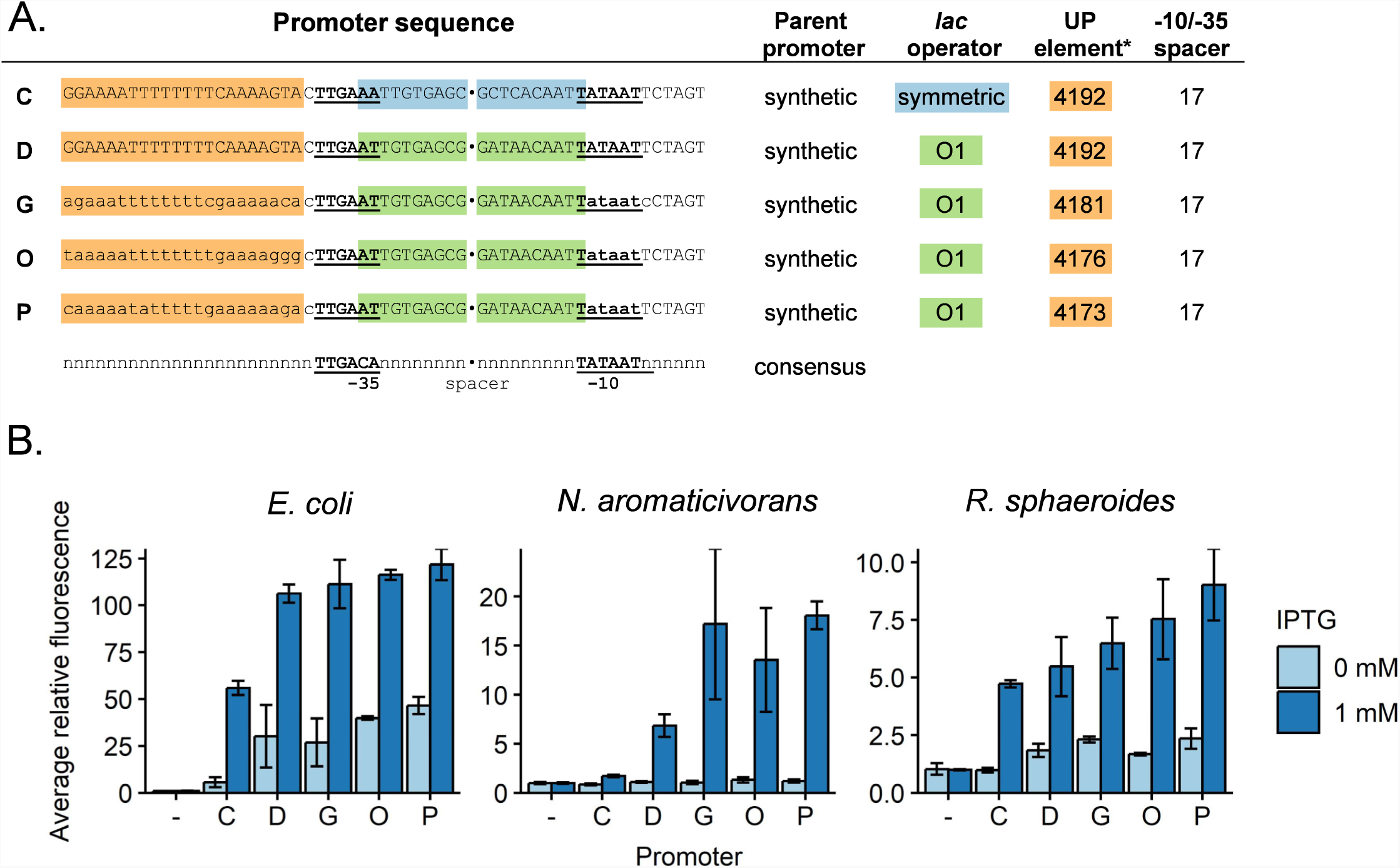
Selected high-activity promoters in *N. aromaticivorans* and *R. sphaeroides*. A: Sequences of five promoters that exhibit the most activity in both organisms. Promoter C includes O1 *lac* operators, while the four others include symmetric *lac* operators. The most sequence differences between the high activity promoters are bases in the UP element. *UP elements correspond to those in Estrem *et al* 1998 (48). B: Activity of each promoter in *E. coli*, *N. aromaticivorans*, and *R. sphaeroides*. A linear scale is used for the y axis. Summary statistics can be found in table S15.

### Programmable gene expression reduction in *N. aromaticivorans* and *R. sphaeroides* with CRISPRi

A specific and significant reduction of gene expression at the transcriptional level can be achieved in many organisms using CRISPRi. CRISPRi utilizes a catalytically inactive Cas9 (dCas9) and gene-targeting single guide RNA (sgRNA) to physically block transcription, reducing gene transcript levels (52). To establish and test the utility of CRISPRi in *N. aromaticivorans* and *R. sphaeroides*, we began by testing the function of a Mobile-CRISPRi system that we developed which inserts in *att*_Tn*7*_ and has been shown to be effective in diverse bacteria (23, 34) (Fig. 4A).

**Figure 4:**
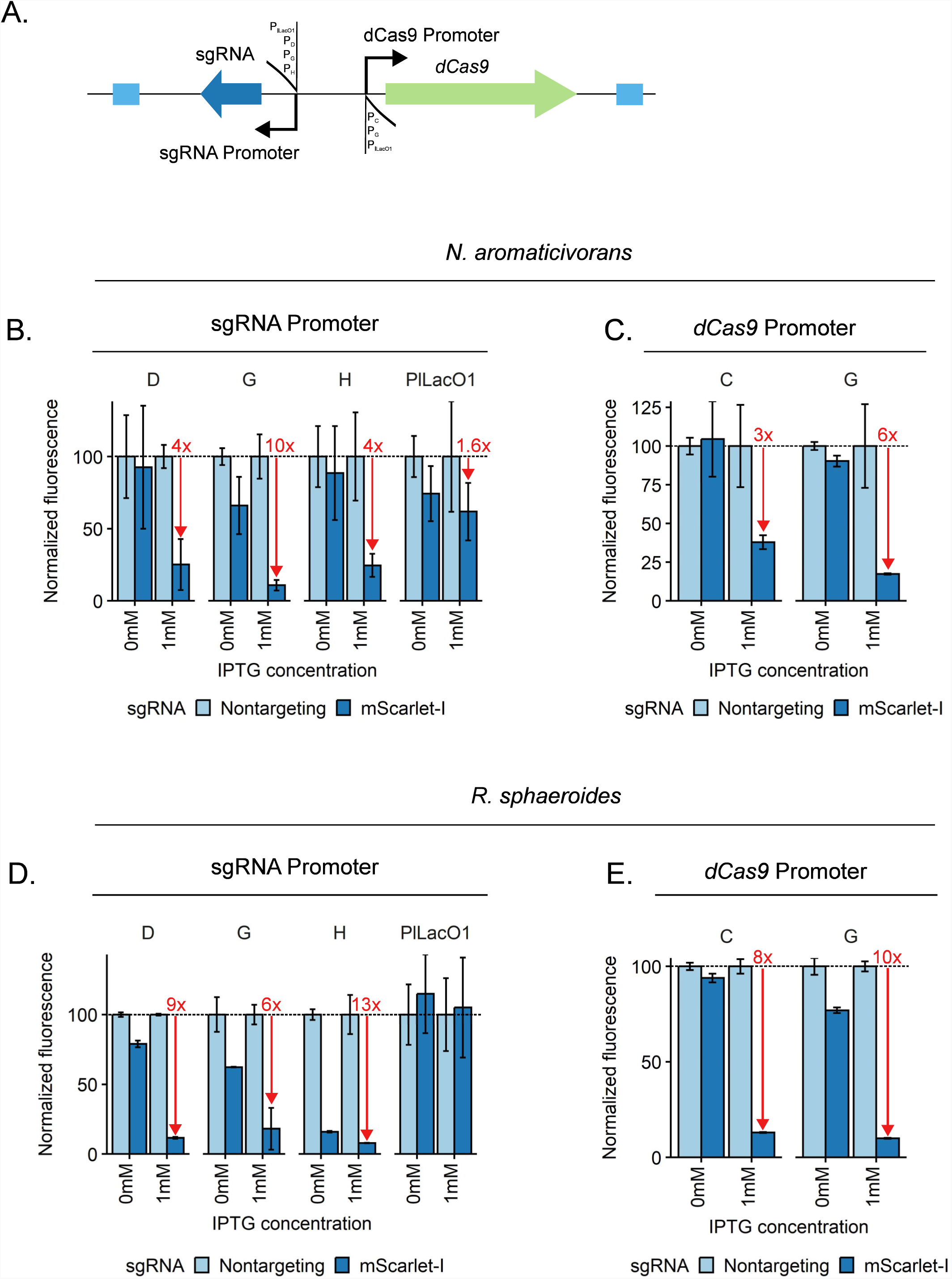
Optimization of Mobile-CRISPRi for use in *N. aromaticivorans* and *R. sphaeroides*. A: Schematic of Mobile-CRISPRi system with varied sgRNA and dCas9 promoters. BD: Use of different promoters to drive sgRNA expression to control *mScarlet-I* expression in *N. aromaticivorans* and *R. sphaeroides*, respectively. All constructs use *Spy-*dCas9 expressed from a P_lLacO1_ promoter. CE: Use of promoters C and G to drive *Spy*-dCas9 expression to control *mScarlet-I* expression in *N. aromaticivorans* and *R. sphaeroides,* respectively. Constructs use promoter D for sgRNA expression. NTC=Non-targeting control. Data normalization: For a given MCi vector, in the given condition, fluorescence per OD600 values for all independent replicates were normalized to the mean fluorescence per OD600 value for the strain encoding non-targeting control sgRNA, which was assigned to be a value of 100% expression. Summary statistics can be found in tables S16-S19.

To test CRISPRi activity, we used a constitutively expressed *mScarlet-I* reporter and either *mScarlet-I*-targeting sgRNAs or non-targeting controls. We also used some of the previously described synthetic promoters that have activity in *N. aromaticivorans* and *R. sphaeroides* to tune expression of the sgRNA (Fig. 4 BD; P_D_, P_G_, and P_H_) and the dCas9 protein (Fig. 4CE; P_C_ and P_G_).The initial Mobile-CRISPRi construct used the P_L*lacO*-1_ promoter to express both dCas9 and sgRNAs (34); use of this construct resulted in negligible reduction in reporter gene activity in both *N. aromaticivorans* and *R. sphaeroides* (Fig. 4BD, Fig S2). However, we found that use of P_D_ or P_H_ resulted in an ∼4-fold reduction in reporter gene activity in *N. aromaticivorans*, while the use of P_G_ increased the reduction in reporter gene activity to ∼10-fold (Fig. 4C). In addition, we found that in *R. sphaeroides*, use of P_D_, P_G_, or P_H_ to control sgRNA expression reduced reporter gene activity 9-, 6-, and 13-fold respectively (Fig. 4D). By testing several combinations of synthetic promoters we found that combining the use of P_D_ for sgRNA transcription with P_C_ or P_G_ for transcription of the *dcas9* gene led to a larger decrease in reporter gene expression in both organisms (Fig. 4CE).The combination of P_D_ for sgRNA transcription with P_C_ or P_G_ for *dcas9* expression yielded similar reductions in reporter gene activity in *R. sphaeroides* (8- and 10-fold, respectively, as compared to 9-fold, initially; Fig. 4E).

To test the ability of CRISPRi to inhibit expression of native genes in *N. aromaticivorans* and *R. sphaeroides*, we targeted the essential *murC* gene (13) that has been shown to be highly sensitive to inhibition in other Gram-negative bacteria (53). In these experiments we tested the P_D_-driven sgRNA and P_C_- or P_G_-dependent dCas9 constructs that showed effective knockdown when using *mScarlet-I* as a reporter (see above). To measure the physiological effects of *murC* knockdown, we grew cells in the absence of induction prior to harvesting and spotting a 10-fold serial dilution series of each strain onto media including or lacking 1mM IPTG (Fig. 5AB). In *N. aromaticivorans*, the P_G_-dependent dCas9 showed the best performance, yielding an ∼10,000x reduction in viability consistent across all replicates (Fig. 5A, compare these results to those obtained using P_C_-dependent dCas9 in Fig. S3A, which shows lower and more variable knockdown). In *R. sphaeroides*, the P_C_ and P_G_-dependent dCas9 constructs performed similarly, with ∼1,000-10,000x reduction in viable colonies in both strains (Fig. 5B, Fig. S4B). In both organisms, we observed no loss of viable colonies in the absence of IPTG, suggesting that expression of CRISPRi components was highly dependent on the addition of IPTG (Fig. 5, Fig. S3). Thus, we conclude that these Mobile-CRISPRi systems for *N. aromaticivorans* and *R. sphaeroides* are effective, highly regulated, and capable of targeting endogenous essential genes in both organisms.

**Figure 5:**
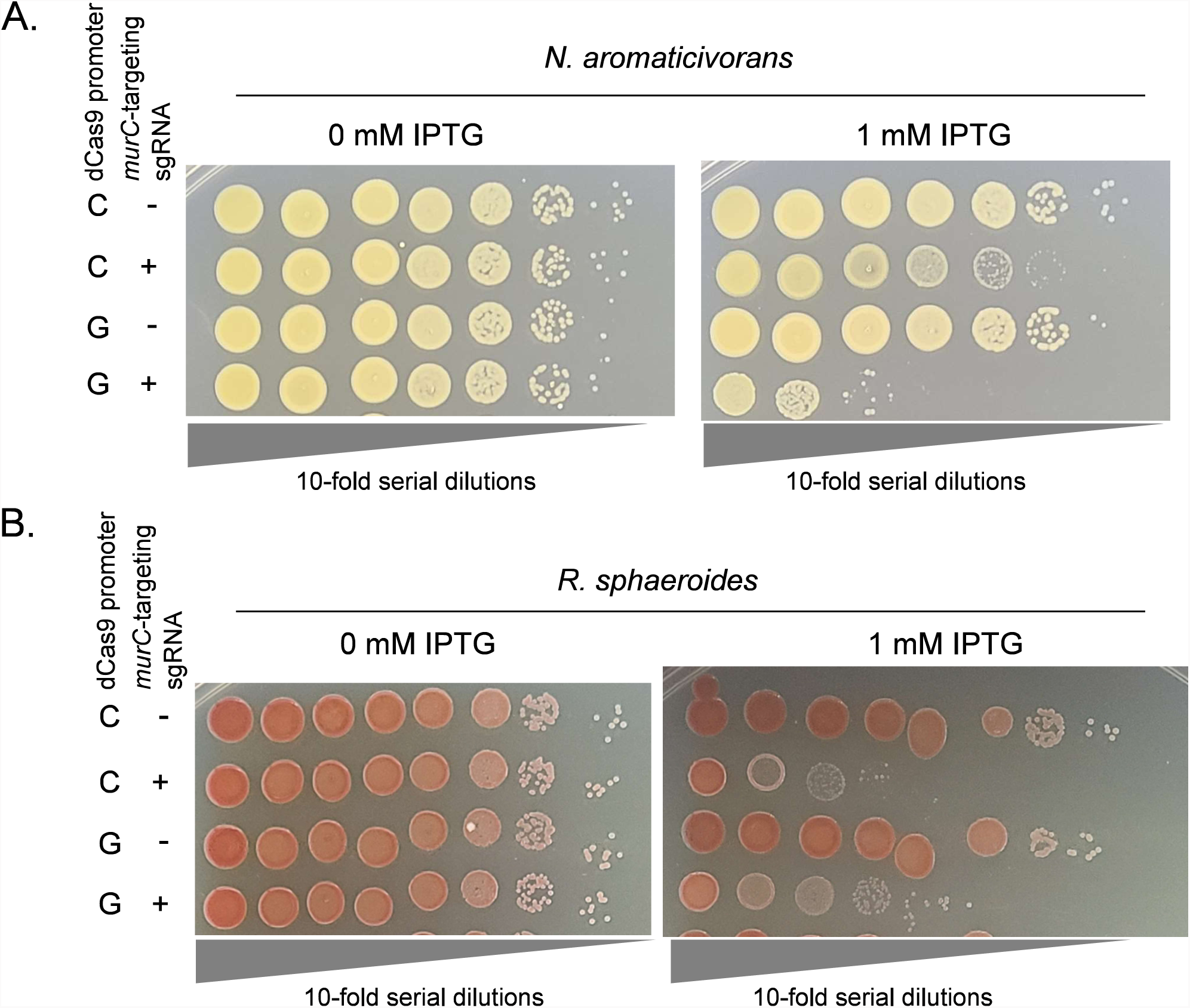
Controlled expression of an essential gene with Mobile-CRISPRi in *N. aromaticivorans* and *R. sphaeroides*. A: 10-fold serial dilutions of *N. aromaticivorans* with MCi constructs with a sgRNA targeting the essential gene *murC* or a non-targeting control sgRNA. *N. aromaticivorans* cells were normalized to an OD600 of 10 prior to serial dilution. Cells were grown on rich media (464a) in the presence or absence of 1mM IPTG. B: 10-fold serial dilutions of *R. sphaeroides* with MCi constructs with a sgRNA targeting the essential gene *murC* or a non-targeting control sgRNA. *R. sphaeroides* cells were normalized to an OD600 of 10 prior to serial dilution. Cells were grown on rich media (LB) in the presence or absence of 1mM IPTG. Figure S3 shows additional biological replicates of each.

### Production and control of an engineered natural product in *N. aromaticivorans*

*N. aromaticivorans* is being developed as an industrial bacterial chassis for the conversion of plant materials into low volume, high value bioproducts (2, 3, 14, 54). We recently demonstrated that the carotenoid astaxanthin can be produced by *N. aromaticivorans* by replacing the genomic copy of the native gene *crtG* with *crtW* from *Sphingomonas astaxanthinifaciens* (Δ*crtG*::*crtW^+^)* (55). To demonstrate the utility of the Tn*7* insertion and inducible promoters described in this work to control the levels of an engineered pathway, we sought to build a strain of *N. aromaticivorans* capable of producing astaxanthin by expression the exogenous *crtW* at the *att*_Tn*7*_ site. We generated and tested several strains in a comparable Δ*crtG* background: two with IPTG-inducible promoters (P_D_ and P_G_) and one with a constitutive promoter (P_H_) driving *crtW* expression. All three strains accumulated astaxanthin in amounts comparable to the original *crtW*-expressing strain when grown in standard laboratory media (Fig. 6A).

**Figure 6:**
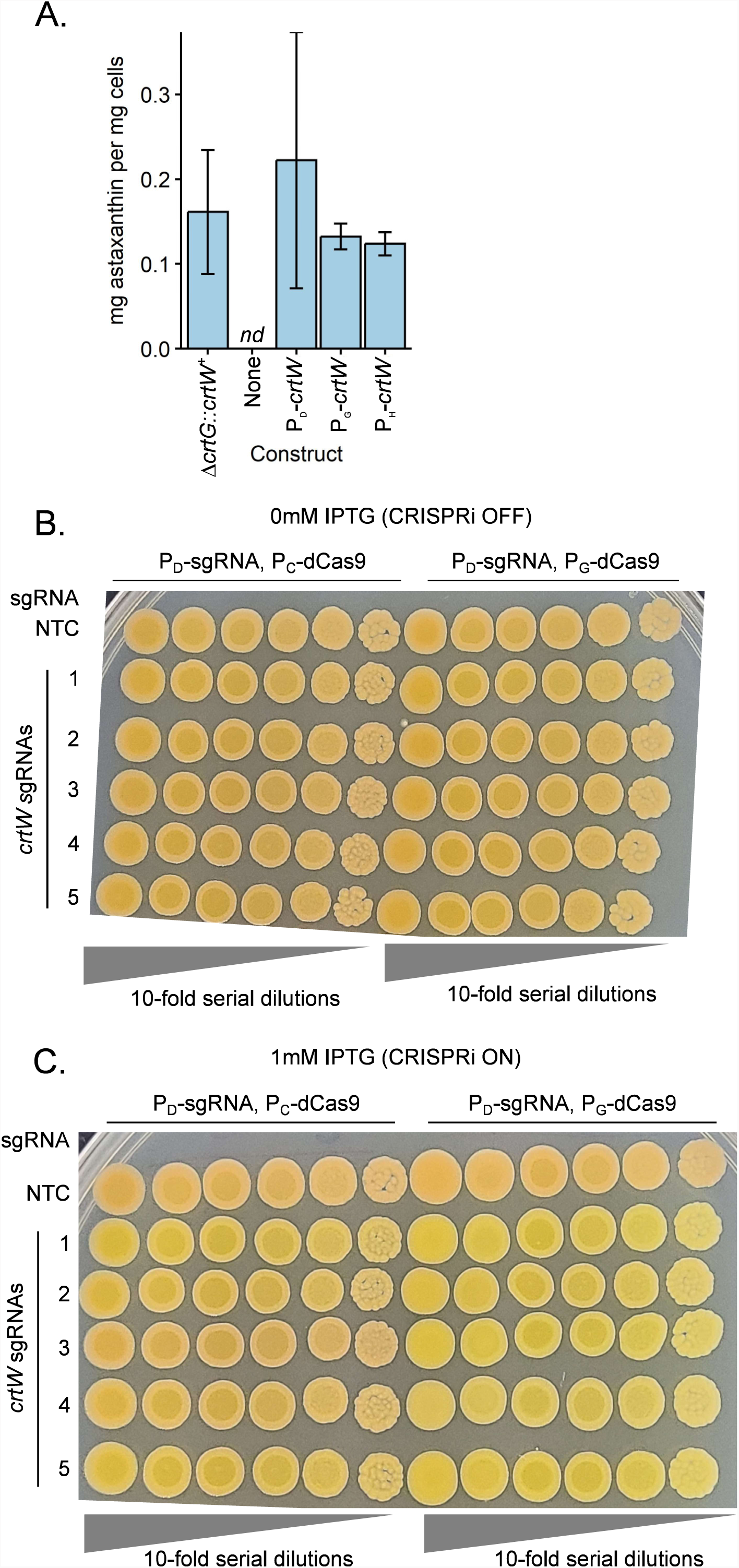
Expression control of the exogenous *crtW* gene in *N. aromaticivorans*. A: Expression of *crtW* from the att::Tn*7* site in Δ*crtG N. aromaticivorans* produces comparable quantities of astaxanthin as a previously-engineered expression strain (Δ*crtG*::*crtW^+^*) (55). There is no difference in the amount of astaxanthin produced among all *crtW* strains; astaxanthin was not detectable in cells lacking this foreign gene (Δ*crtG*, Tn*7* construct “None”). B: 10-fold serial dilutions of Δ*crtG*::*crtW^+^* cells with an MCi system at the att::Tn*7* site. Cells were plated on rich media (464a) lacking (B) or supplemented with (C) 1mM IPTG. n.d. = Not detected. Summary statistics can be found in table S20.

To further test the utility of the Tn*7* insertion system we sought to use CRISPRi to control the expression of this foreign *crtW* gene. Using the Δ*crtG*::*crtW^+^* strain, we inserted Tn*7*-based CRISPRi constructs that carried either a nontargeting control guide or a one targeting *crtW*. We also tested constructs in which P_C_ or P_G_ was used to control transcription of the *dcas9* gene. We know that CrtW expression results in orange colonies due to astaxanthin accumulation, so we used colony pigmentation as a visible reporter for changes in *crtW* expression. In the absence of CRISPRi induction, colonies from all strains were orange, suggesting they accumulated astaxanthin (Fig. 6C). In contrast, in the presence of IPTG to induce transcription of CRISPRi, some strains with P_C_-*dcas9* (guides 1, 2, and 5) and all strains with P_D_-*dcas9* became yellow (characteristic of Δ*crtG N. aromaticivorans*), indicating a reduction in *crtW* expression. Based on colony color, when paired with P_C_-*dcas9*, *crtW* guides 3 and 4 were visually less efficacious than guides 1, 2, and 5. Overall, the P_D_-*sgRNA*, P_G_-*dcas9* construct produced a more consistent knockdown for the *crtW* gene when expressed in *N. aromaticivorans*. Taken together, we find that combining use of Tn*7*, select synthetic promoters, and CRISPRi can be used to effectively build and modulate gene expression of engineered constructs in *N. aromaticivorans*.

## DISCUSSION

There is a need for facile tools for genomic engineering and editing to dissect the lifestyle of Alphaproteobacteria and to remove barriers in efficiently constructing strains for production of bioproducts from abundant renewable raw materials including deconstructed lignocellulosic biomass. The suite of tools presented here for *N. aromaticivorans* and *R. sphaeroides* advance our ability to engineer pathways and will accelerate strain optimization. We show that our platform of Tn*7*, synthetic promoters, and CRISPRi effectively deliver and modulate gene expression of native genes and engineered constructs in *N. aromaticivorans* and *R. sphaeroides*. These genetic tools can be applied to both gene function discovery and manipulating metabolic pathways, providing the means to identify engineering targets and further manipulate gene expression to obtain new knowledge about the biology of these microbes, possibly other Alphaproteobacteria as well as other members of the bacterial phylogeny.

Our use and optimization of Tn*7* transposition in *N. aromaticivorans* and *R. sphaeroides* provides a rapid, straightforward approach for genomic integration of single genes or entire pathways. With the efficiencies observed here, constructing pathway libraries on the order of thousands of permuted pathways should be feasible; this will enable high-throughput screening of gene product and pathway function directly in *N. aromaticivorans* and *R. sphaeroides*. Although the cargo capacity of Tn*7* is unknown, the original Tn*7* element contains ∼14 kb of DNA (56) and Tn*7*-like elements associated with CRISPR systems (CAST) range in size from 22-120 kb (57); therefore, we propose that the size of the inserted DNA sequence is unlikely to be a limiting factor for Tn*7* integration. The Tn*7* attachment site can also be used for complementation analysis to verify that the function of a specific gene is central to an observed phenotype, allowing the investigator to avoid well-documented *cis*-acting affects, such as polarity, onto downstream genes as previously shown in other bacteria (58).

Our findings that Tn*7* transposition efficiency can be impacted by mating strategy and, presumably, transposase gene expression both sheds light on past difficulties in using this transposon in some species and opens additional avenues for increasing the recovery of transconjugants for library-scale studies. The inclusion of an additional *E. coli* donor strain containing a conjugal helper plasmid increased Tn*7* transconjugant recovery by an unknown mechanism. Although all *E. coli* strains present in the mating expressed conjugation machinery, it is possible that multicopy expression of the conjugation machinery drove recovery of additional transconjugants. We also observed similar behavior in *Acinetobacter baumannii* conjugations (59), indicating that the phenomenon could be broadly applicable. Expression of Tn*7* transposase genes in *R. sphaeroides* substantially increased recovery of transconjugants, suggesting that transposase gene expression was limiting. Our findings that Tn*7* transposition efficiency can be impacted by mating strategy and, presumably, transposase gene expression both sheds light on past difficulties in using this transposon in some species and opens additional avenues for increasing the recovery of transconjugants for library-scale studies.

Our identification of synthetic, inducible promoters for *N. aromaticivorans* and *R. sphaeroides* shows that some promoters built on the *E. coli* σ^70^ consensus can drive high-level expression of native and foreign genes in Alphaproteobacteria, although not all promoters that were active in *E. coli* were also active in *N. aromaticivorans* and *R. sphaeroides*. It is possible that the reported inactivity of the *E. coli lac* promoter in *R. sphaeroides* (26) may be due to suboptimal −10 or −35 sequences, a wide spacer sequence (*lac* has an 18 bp spacer, rather than the more standard 17 bp), or some combination of elements. Interestingly, P_A_—a variant of the *lacUV5* promoter with an 18 bp spacer—shows higher relative activity in *N. aromaticivorans* than *R. sphaeroides*, suggesting there can be organism-specific differences in promoter sequence requirements between individual species. All of our promoters contained a “T” base at the −7 position of the −10 element; therefore, we were unable to test if a substitution of this base would be tolerated in *N. aromaticivorans* and *R. sphaeroides* as recent work has shown for the ribosomal RNA promoter of *R. sphaeroides* (41). Retaining a “T” at the −7 position may still be advantageous in Alphaproteobacteria to avoid dependence on *trans* acting factors such as the CarD transcription factor (33, 41, 60).

We predict that the use of CRISPRi in *N. aromaticivorans* and *R. sphaeroides* will provide a powerful new genetic tool to complement existing homology-mediated genome modification and Tn-seq in these and other Alphaproteobacteria (2, 3, 14). For example, CRISPRi will permit genome-scale interrogation of essential genes in *N. aromaticivorans* and *R. sphaeroides* as we have recently demonstrated in *Z. mobilis* (24). Also, Mismatch-CRISPRi—the use of sgRNAs that are deliberately modified to imperfectly match target DNA—will allow partial knockdowns of essential genes (53) that may be relevant for directing metabolic flux toward bioproducts. Combining CRISPRi functional genomics with metabolic models (61) may further accelerate rational engineering to increase bioproduct yields. We expect that these approaches will be useful in dissecting metabolic and regulatory networks in Alphaproteobacteria and in testing hypotheses for engineering strains for bioproduct synthesis. Together, these advancements provide the foundation from which to increase our understanding of microbial activities and to propel the bioeconomy.

## METHODS

### Strains and growth conditions

Table 1 lists all strains used in this study. *E. coli* strains were grown in LB (10 g tryptone, 5 g yeast extract, 5 g NaCl per liter; BD 240230) aerobically, either at 37°C in a flask with shaking at 250 rpm, in a culture tube on a roller drum, in a 96 deep-well plate with shaking at 900 rpm or at 30°C in a flask or a culture tube with shaking at 250rpm. *R. sphaeroides* strains were grown in Sistrom’s Minimal Media (SIS) (62) or LB at 30°C in a flask or a culture tube with shaking at 250rpm. *N. aromaticivorans* strains were grown in 464a (5g tryptone, 5g yeast extract, 1g glucose per liter) (DSMZ) or SIS supplemented with 1% glucose.

For growth on plates, 15g/L agar was added to the appropriate medium. When necessary, antibiotics were used at the following concentrations: *E. coli*: 100 μg/ml ampicillin (amp) or 30 μg/ml kanamycin (kan). *R. sphaeroides*: 25 μg/ml spectinomycin (spec), 25 μg/ml kan. *N. aromaticivorans*: 30 μg/ml kan, 15μg/mL gentamycin (gent). 1 mM Isopropyl β-D-1-thiogalactopyranoside (IPTG) was added where indicated.

All strains were preserved in 15-25% glycerol and frozen at −80°C.

### Conjugation and Tn*7* transposition

#### Growth of strains

##### Donor strains

The transposase donor strain sJMP2591 or sJMP11111, helper plasmid strain sJMP11115, and transposon donor strain (Table 1) were grown on LB agar supplemented with DAP (300μM) and the appropriate antibiotic and grown 16-20 hours at 37°C. Donor cultures were started from a single colony in 10mL LB broth supplemented with DAP and amp and grown with shaking (250rpm) at 30°C for 16-20 hours. Alternately, cells were scraped from the primary streak after overnight growth and resuspended in LB.

##### *E. coli* recipient

DH10B F- strain sJMP3272 was streaked out on LB agar and grown overnight at 37°C. Recipient cultures were started from a single colony in 10mL LB broth and grown with shaking (250rpm) at 30°C for 16-20 hours. Alternately, scrapes of each the donor and recipient strains were mixed on solid media, grown 2-8 hours at 37°C, then streaked for single colony isolates on selective media.

##### R. sphaeroides recipient

Mating strategy used depends on recipient strain.

##### sJMP8063

Strain sJMP8063 containing the transposase-expressing plasmid pJMP8045 was streaked out on SIS media supplemented with 25μg/mL spec and grown for one day at 30°C. A pre-culture from a scrape of cells from the primary streak was inoculated in one milliliter of SIS supplemented with 25μg/mL spec and grown with shaking (250rpm) at 30°C for 18-22 hours. This 1mL pre-culture was then added to 10mL SIS supplemented with 25μg/mL spec and grown with shaking (250rpm) at 30°C for another 18-22 hours, then used in conjugation.

##### sJMP8005

Strain sJMP8005 is wild type *R. sphaeroides* and a quad-parental mating scheme was used for mating of Tn*7* constructs. Strain sJMP11111 or sJMP2591 was used as the *E. coli* transposase donor, strain sJMP11115 as the conjugal helper plasmid donor, and the appropriate transposon donor was selected; see table 1.

### N. aromaticivorans recipient

Promoter-mScarlet-I and Fig. S3 CRISPRi constructs: Strain sJMP8004 was streaked on 464a plates and incubated at 30°C for two days. A 10mL 464a culture was inoculated from a single colony and grown for 16-20 hours at 30°C in a 125mL baffled flask with shaking at 250rpm. Cells were harvested and resuspended to an OD600 of 3. 1mL of cells was used in each mating, with 200μl of each *E. coli* transposase and transposon donor, also normalized to an OD600 of 3. Cells were concentrated by centrifugation at 4000xg, supernatant was decanted, and cells were resuspended in residual volume, spotted on plates, and allowed to mate overnight. Mating spots were then scraped into 1mL 464a media, resuspended, serially diluted, and plated.

Figure 1 mating efficiency experiments, all other CRISPRi constructs and *crtW* strains: Strain sJMP11093 was patched onto 464a plates directly from the freezer and incubated at 30°C for ∼24 hours. Cells were scraped into 464a media and normalized to an OD600 of 3. 1mL of cells was combined with *E. coli* donor cells: 200μl transposase donor at OD600 3, 400μl transposon donor at OD600 3, and 400μl conjugal helper at OD600 3. Cells were concentrated by centrifugation at 4000xg, supernatant was decanted, and cells were resuspended in residual volume, spotted on plates, and allowed to mate for four hours at 30°C. Mating spots were then scraped into 1mL 464a media, resuspended, serially diluted, and plated.

### Cloning

#### Plasmid preparation

Plasmids were prepared from *E. coli* strains with one of the following kits: GeneJet plasmid miniprep kit (catalog number K0503; Thermo Scientific). QIAprep Spin Miniprep Kit (catalog number 27104; Qiagen), or the PureLink HiPure Plasmid Midiprep kit (catalog number K210005; Invitrogen).

#### Preparation of competent E. coli cells

Competent cells of strains sJMP3053 (*pir*+ cloning strain) and sJMP3257 (*pir*+, *dap*- mating strain) were prepared as previously described (34). Briefly, *E. coli* cells were grown in LB (supplemented with DAP when appropriate) at 37°C to early log phase (OD600 ∼0.3). Cells were immediately chilled on ice, harvested by centrifugation in a swinging-bucket rotor, then washed 3x in cold 5% glycerol. Aliquots of cells suspended in 15% glycerol were stored at −80°C for later use.

#### Transformation

*E. coli* cells were electroporated with the Bio-Rad Gene Pulser Xcell on the EC1 setting in 0.1cm cuvettes. Cells were recovered in 800μl LB media for one hour at 37°C prior to plating on appropriate antibiotic, adding DAP when necessary.

#### Sequence validation

Sanger sequencing of select plasmid regions or select inserts was performed by Functional Biosciences (Madison, WI). Whole-plasmid long-read sequencing was performed by Plasmidsaurus (Eugene, OR) using Oxford nanopore technology.

#### pAlphabet-mScarlet-I constructs

Parent vector pJMP8602 was digested with restriction enzymes PacI (NEB) and SpeI (NEB). Complementary oligonucleotides with overhangs compatible with the PacI/SpeI-digestd pJMP8602 sticky ends were annealed by combining equimolar amounts (2μM each) in 1X Cutsmart buffer (NEB), heating to 95°C for 10 minutes, then allowing to gradually cool to room temperature. Annealed oligos were diluted 1:20and 2ul of annealed oligonucleotides was ligated into the digested pJMP8602 with T4 DNA ligase (NEB 0491) for either two hours at room temperature or 16 hours at 16°C. Two microliters of the ligation reaction were directly transformed into 50μl electrocompetent sJMP3053 cells and plated on appropriate selective medium. Clones were single-colony purified prior to confirming plasmid construction and preservation at −80°C.

#### CRISPRi constructs

Parent Mobile-CRISPRi constructs were built as indicated in Table 2. sgRNA spacer sequences were cloned into the BsaI site of each relevant Mobile-CRISPRi vector as previously described (24, 34). Briefly, a 20-nucleotide sequence proximal to an NGG PAM sequence was identified, complementary to the gene of interest. 4-nucleotide ends were added to oligos encoding each strand such that upon annealing, the pair will have complementary overlap to the BsaI cut ends of the Mobile-CRISPRi vector. Sequences of oligonucleotides used to build CRISPRi constructs can be found in Table 3.

### Tn*7* insertion stability

#### N. aromaticivorans

Starter cultures were grown in 464a supplemented with kan overnight at 30°C with 250rpm shaking. One mL of culture was harvested by centrifugation, supernatant decanted, and resuspended in 1mL media lacking antibiotics. Cells were diluted 1:1000 into fresh media and grown 24h. The process of diluting 1:1000 and growing 24h was repeated a total of five times for approximately 50 generations of growth in the absence of selection. Cells were serially diluted and plated on media lacking antibiotics, and 150 colonies per biological replicate were patched onto 464a and 464a+kan to determine the fraction that retained the Tn7 insertion.

#### R. sphaeroides

Samples were prepared as for *N. aromaticivorans* but using SIS media and an alternative dilution and timing process to account for the higher doubling time. The growth and dilution protocol was performed a total of six times, one of which was a 1:100 dilution and the remainder of which were 1:1000 dilutions. The total number of generations of growth in the absence of selection was approximately 50-55 generations.

### pTNS++ plasmid curing

A strain carrying pTNS++ was streaked out from the freezer onto rich media lacking antibiotics. Single colonies were inoculated in liquid media lacking antibiotics and grown 24 hours, then serially diluted, and plated on media lacking antibiotics. Plates were incubated for 3 days to obtain colonies, which were patched onto selective and nonselective plates.

### Growth curves

Three biological replicates per genotype were grown to saturation in liquid media overnight, in the media in which the growth curve would be completed. Samples were diluted 1:1000 (*N. aromaticivorans* complete and minimal media, *R. sphaeroides* complete media) or 1:100 (*R. sphaeroides* minimal media) into a total volume of 200μl in a clear, flat-bottomed 96-well plate (Corning). The following concentrations were used for each additive: 1mM IPTG, 2% DMSO, 2mM Theophylline (2% DMSO). Growth curves were performed in a Tecan Sunrise or Tecan Infinite 200 Pro Mplex at 30°C, measuring OD600 every 15 minutes.

Data from the growth curves were analyzed in R with the growthcurver package (63).

### Fluorescence assays

Starter cultures were prepared either by growing in liquid medium overnight from single-colony isolates, or by patching single-colony isolates onto agar plates and the resulting growth scraped into liquid and OD600-normalized the next day.

Starter cultures were diluted and inoculated into 96-deepwell plates containing the appropriate media for the species analyzed, with or without 1mM IPTG, 2mM Theophylline, or 2% DMSO. Plates were incubated at 30°C on a plate shaker (Benchmark Orbi-shaker MP) set to 1000rpm until cells had grown to saturation. 200μl of each culture was transferred to a black, clear-bottom plate for analysis in a Tecan Mplex Infinite platereader.

### Spot plate assays

Triplicate single colony isolates of *N. aromaticivorans* or *R. sphaeroides* were patched onto appropriate media lacking inducer and grown overnight at 30°C. Cells were scraped off plate and resuspended in 1mL appropriate media (464a or LB) and normalized to an OD600 of 10. 10-fold serial dilutions of each strain were prepared in 96-well plates, and 5μl of each was spotted onto 15cm petri plates containing the appropriate media either with or without 1mM IPTG. Plates were grown at 30°C and imaged over 2-5 days in a lightbox with a Samsung Galaxy S20+ phone camera.

### Quantification of astaxanthin

Astaxanthin levels were quantified as previously described in Hall *et al* 2023 (55).

#### Cell growth

Starter cultures were made by inoculating single colonies of *N. aromaticivorans* isolates in 2mL 464a and growing for approximately 15 hours at 30°C (not to saturation). Samples were diluted 1:50 into a total volume of 25mL 464a, and 1mM IPTG was added to appropriate samples. Cells were grown 9h at 30°C with shaking at 250rpm. Two 10mL aliquots were harvested from each culture by centrifugation and processed as Follows. 1: Supernatant was decanted and pellet was allowed to dry in chemical fume hood to calculate dry cell weight. 2: Supernatant was decanted and pellets were flash frozen in liquid nitrogen and stored at −80°C until preparation of lipophilic extracts.

#### Preparation of lipophilic extracts

Cell pellets were resuspended in 200 μL water, then transferred into a 15 mL centrifuge tube. 4 mL extraction solvent (7:2 acetone:methanol solution) was added and the samples were mixed by pipetting. The tube was centrifuged (10,000 × g for 20 min), then the supernatant was transferred to a new 15 mL tube. The pelleted cells were extracted a second time, adding 100 μL water for resuspension followed by 4 mL extraction solvent. After centrifugation, the supernatants from both extractions were combined. The combined supernatants were partially dried under a stream of N2 (to a final volume of ∼1 – 3 mL) to concentrate materials before analysis by HPLC.

#### HPLC identification and quantification of astaxanthin

Acetone:methanol lipophilic extracts were analyzed via reverse-phase HPLC using a Kinetex 2.6 μm PS C18 100 Å (150 x 2.1 mm) column (Phenomenex; Torrance, CA) attached to a Shimadzu Nexera XR HPLC system. The mobile phase was a binary gradient of Solvent A (70% acetonitrile/30% water) and Solvent B (70% acetonitrile/30% isopropanol) flowing at 0.45 mL/min. Absorbance was measured between 200 and 600 nm using a Shimadzu SPD- M20A photodiode array detector. An astaxanthin commercial standard (Sigma-Aldrich) was used to identify and quantify astaxanthin in the extracts. Astaxanthin eluted at a retention time of 12.6 min.

## ACKNOWLEDGEMENTS

This material is based upon work supported by the Great Lakes Bioenergy Research Center, U.S. Department of Energy, Office of Science, Office of Biological and Environmental Research under Award Number DE-SC0018409. B. W. H. was partially supported by NIH Training Grant #5T32GM007133.

---

**Figure S1:**
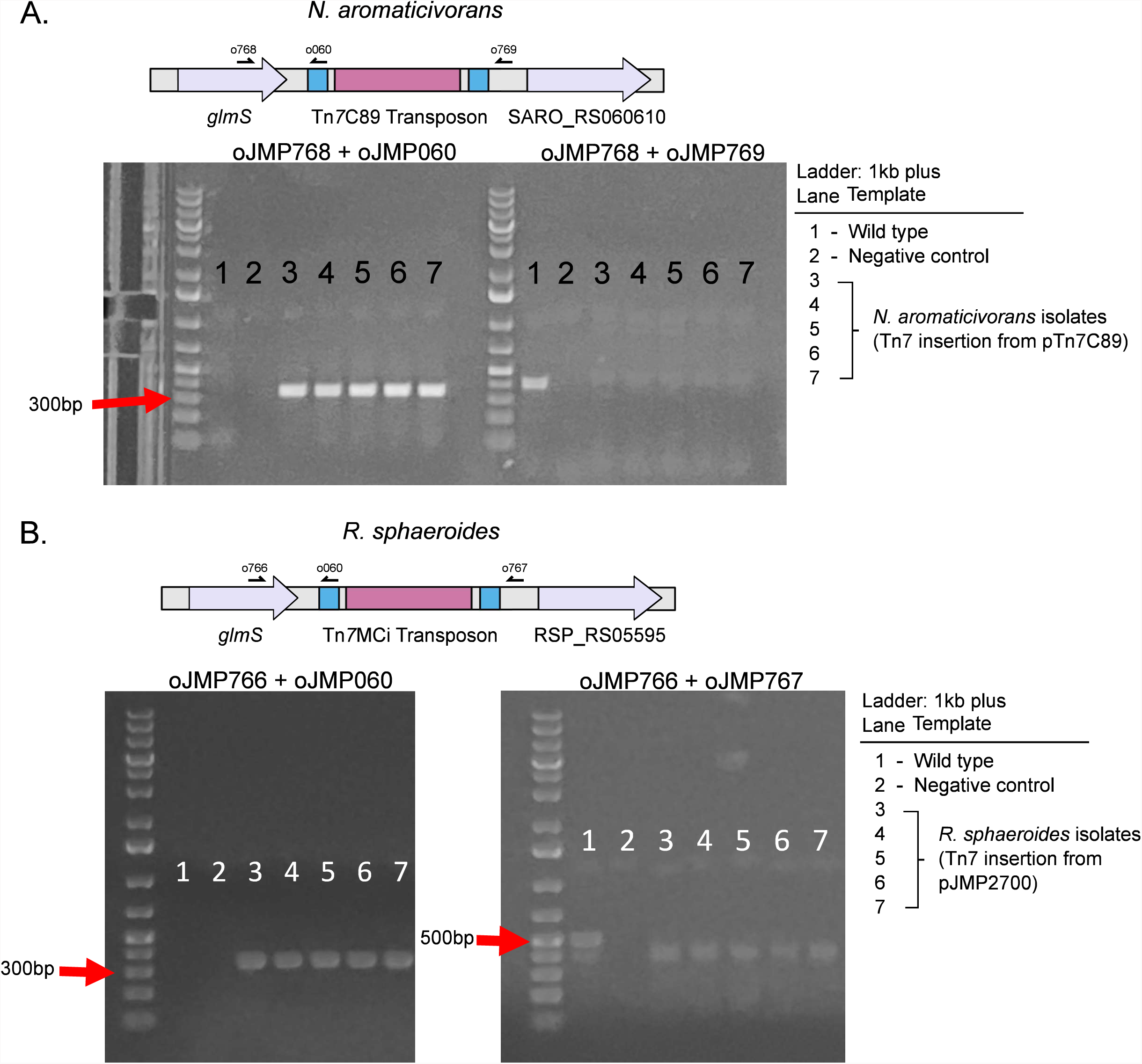
Confirmation of att::Tn*7* integration. A: Schematic of *glmS* genetic region with and without att::Tn7 integration and locations of validation primers. B: *N. aromaticivorans* clones. C: *R. sphaeroides* clones.

**Figure S2:**
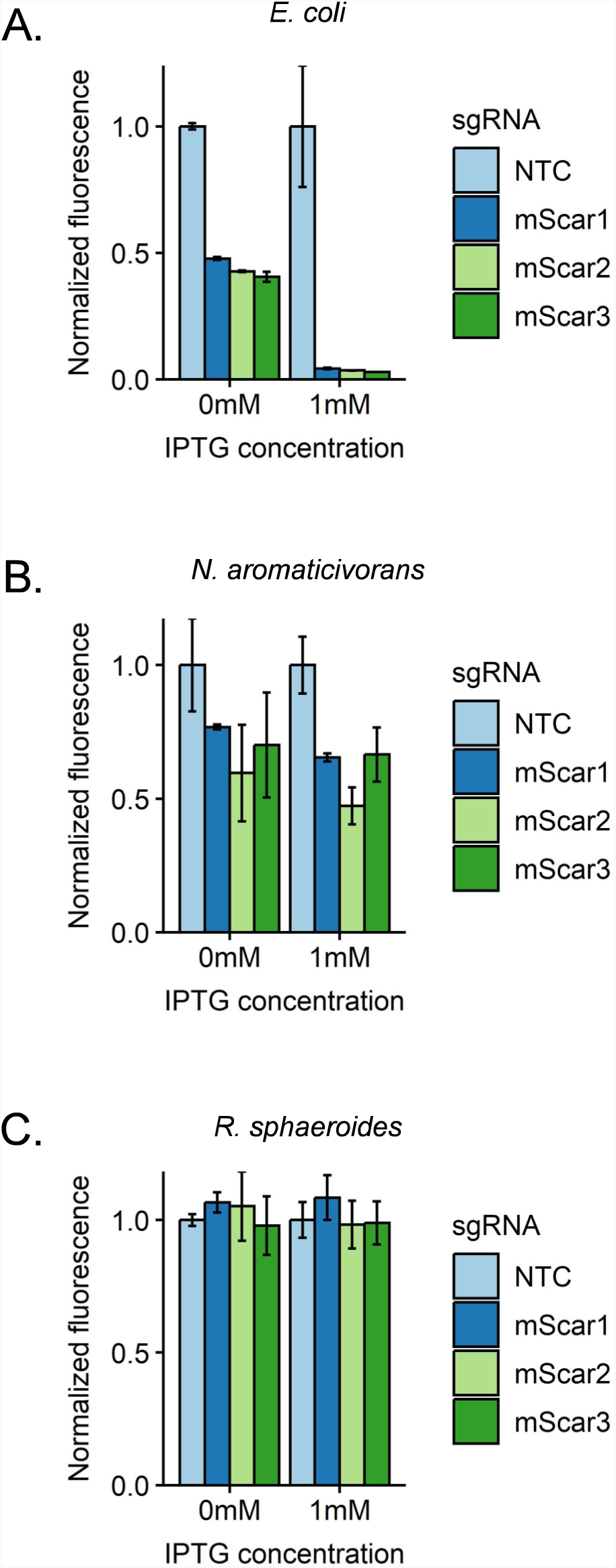
CRISPRi knockdown of *mScarlet-I* with initial Mobile-CRISPRi constructs. Constructs with both sgRNA and dCas9 under the control of PlLacO1 were tested with three different targeting sgRNAs in *E. coli* (A), *N. aromaticivorans* (B), and *R. sphaeroides* (C). NTC=Non-targeting control strain.

**Figure S3:**
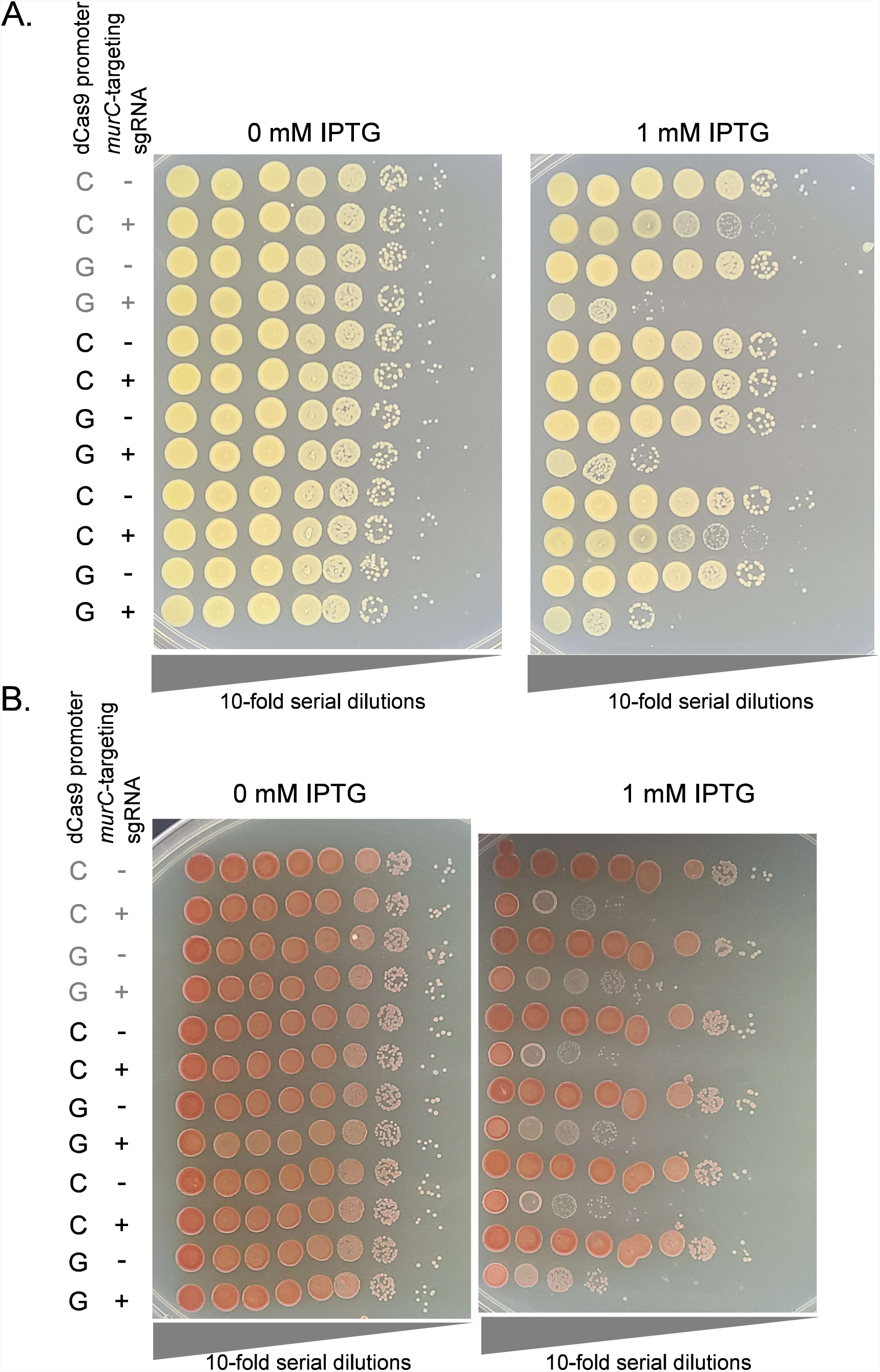
Expanded data for *murC* CRISPRi in *N. aromaticivorans* and *R. sphaeroides*. A: 10-fold serial dilutions of *N. aromaticivorans* with MCi constructs with a sgRNA targeting the essential gene *murC* or a non-targeting control sgRNA. *N. aromaticivorans* cells were normalized to an OD600 of 10 prior to serial dilution. Cells were grown on rich media (464a) in the presence or absence of 1mM IPTG. B: 10-fold serial dilutions of *R. sphaeroides* with MCi constructs with a sgRNA targeting the essential gene *murC* or a non-targeting control sgRNA. *R. sphaeroides* cells were normalized to an OD600 of 10 prior to serial dilution. Cells were grown on rich media (LB) in the presence or absence of 1mM IPTG. Samples presented in the main text are indicated in gray.

